# Remarkable Biosynthetic Diversity in the *Bionectriaceae*: Uncovering the genomic basis of antipathogen bioactivity of cacao fungal endophytes *Clonostachys rosea* and *Waltergamsia zeylanica*

**DOI:** 10.64898/2026.11.16.767117

**Authors:** Librada A. Atencio, Hilda E. Castillo, Masiel Barrios-Jaén, Indira Quintero, Andrea Martínez, Candelario Rodriguez, Marcelino Gutiérrez, Luis C. Mejía

**Affiliations:** División de Biodiversidad, Instituto de Investigaciones Científicas y Servicios de Alta Tecnología, Ciudad del Saber, Ancón, Republica de Panamá; Universidad Tecnológica de Panamá; Estación Científica Coiba, Ciudad del Saber, Ancón, Panamá, República de Panamá; División de Bioprospección y Descubrimiento de Fármacos, Instituto de Investigaciones Científicas y Servicios de Alta Tecnología, Ciudad del Saber, Ancón, Republica de Panamá; Smithsonian Tropical Research Institute, Ancon, Panama, Republic of Panama

**Keywords:** *Bionectriaceae*, fungal endophytes, biosynthetic gene cluster, *Clonostachys rosea*, *Waltergamsia zeylanica*

## Abstract

Cacao (*Theobroma cacao*) production in tropical countries is threatened by fungal diseases caused by *Moniliophthora roreri* and *M. perniciosa*. Two cacao pod endophytes from *Bionectriaceae*, *Clonostachys rosea* TCF417 and *Waltergamsia zeylanica* TCF400, exhibit growth antagonism towards *M. roreri*; however, the genetic basis underlying the bioactivity of these strains has not been addressed. Here, we aimed to identify biosynthetic gene clusters (BGCs) potentially associated with the antifungal bioactivity of these two strains and performed a comparative analysis with 30 publicly available *Bionectriaceae* genomes using BiG-SCAPE. Whole-genome sequencing using Oxford Nanopore and Illumina technologies, resulted in assemblies of 52.9 Mb (*C. rosea* TCF417) and 31 Mb (*W. zeylanica* TCF400). Genome annotation was conducted using Funannotate, and secondary metabolite BGCs were predicted using antiSMASH. The two cacao endophytes harbored 92 BGCs (*C. rosea*: 67; *W. zeylanica*: 25), indicating remarkable biosynthetic diversity. Across all 32 genomes, 1,523 BGC-containing regions were identified, most belonging to T1PKS clusters. Network-based clustering revealed 209 gene cluster families (GCFs), of which 52 corresponded to previously characterized BGCs. Conserved BGCs for antifungal compounds, including sorbicillinoids and verticillins, were identified in cacao endophytes and across *Bionectriaceae*. Additionally, an antifungal assay conducted here demonstrated that verticillin D from *C. rosea* TCF417 exhibited growth inhibition of plant pathogens *Moniliophthora roreri* and *Mycena citricolor*. With about 75% of unassigned GCFs, our results reveal a largely untapped biosynthetic repertoire in the *Bionectriaceae* and identify verticillin D as a candidate contributor to the antagonism of *C. rosea* against plant pathogens.

## Introduction

Global production of cacao (*Theobroma cacao*) is threatened by various diseases. Pathogens such as *Phytophthora* spp. (black pod disease), *Moniliophthora roreri* (frosty pod rot), and *M. perniciosa* (witches broom) are the main causal agents of cacao diseases in Latin America and generates significant declines in this crop production (Marelli et al. 2019; International Cocoa Organization 2025). Long-term sustainable solutions have been proposed for the management of these diseases, including the use of biological control agents and integrated pest control management (Hanada et al. 2010). Endophytic fungi have been proposed as biocontrol agents to combat pathogens that affect cacao based on studies that shows their capacity to control the growth of the most common and economically important pathogens of this crop tree (Mejia et al. 2008; Hanada et al. 2010; ten Hoopen and Krauss 2016; Castillo et al. 2025). Certain members of the fungal family *Bionectriaceae* have been documented to function as biocontrol agents (BCA) of plant pathogens. *Clonostachys rosea* has been extensively documented to provide benefits such as growth promotion and disease control in several crop plants including cacao, and through various mechanisms, including mycoparasitic activity, fungal cell wall degradation, and production of secondary metabolites (Krauss and Soberanis 2001; Mejia et al. 2008; ten Hoopen and Krauss 2016; Sun et al. 2020; Zhao et al. 2023). In addition, *Verruciconidia persicina* (formerly known as *Acremonium persicinum*) has been utilized as a BCA against coconut tar spot and, has been reported to inhibit the germination of *Hemileia vastatrix* for the biological control of coffee leaf rust (Bettiol 1996; Liang et al. 2025; Zhao et al. 2025)

Research on the metabolic capabilities of *Bionectriaceae* has primarily focused in the genus *Clonostachys,* where a wide range of compounds with antifungal activity had been reported, including peptaibiotic metabolites, clonorosein A and B, docosapentaenoic acid, cinnamic acid, among others (Han et al. 2020; Sun et al. 2020; Jaffer et al. 2025). However, little is known about the diversity of secondary metabolites produced by this fungal family. Therefore, there is a need to investigate the biosynthetic potential encoded in the genomes of these fungi. The genes involved in the biosynthesis of fungal secondary metabolites are typically arranged in biosynthetic gene clusters (BGCs), and their architecture has facilitated the development of algorithms to predict BGCs that encode specialized enzymes (synthases and synthetases) in the fungal genome (Keller 2019; Hemmerling and Piel 2022). Generally, BGCs contain complete pathways that facilitate the biosynthesis, assembly, modification, resistance, and regulation of their products (Hemmerling and Piel 2022). Biosynthetic enzymes, such as Polyketide synthases (PKS), non-ribosomal peptide synthetases (NRPS), and terpenes synthases, are of great importance in fungi. PKSs are multi-domain enzymes with multiple functional domains that synthesize polyketides. The metabolites generated have diverse structures and biological activities, and are involved in cell growth and development, environmental adaptation, and pathogenicity in many fungi (Fatema et al. 2018; Keller 2019; Stroe et al. 2024). NRPS are complex, multimodular enzymes that produce peptides via non-ribosomal mechanisms. These enzymes are known for producing compounds with diverse bioactivities, including antifungal and antibacterial properties. Furthermore, they are involved in fungal development and stress tolerance (Niu et al. 2020; Oide and Turgeon 2020; Zhang et al. 2023). Terpene synthases are key enzymes in the biosynthesis of terpenes and terpenoids, which serve as chemical defenses against predators, contribute to cellular functions, and facilitate the establishment of symbiotic relationships (González-Hernández et al. 2023).

As genomic sequencing of fungal strains becomes economically viable, studying the entire genome of biocontrol agent candidates is valuable for identifying and enhancing our understanding of the genomic basis of their bioactivity. Members of *Bionectriaceae* are widely distributed in different environments, mostly terrestrial and freshwater habitats, although some species are also present in marine ecosystems. They exhibit diverse lifestyles, including, coprophilous, herbicolous, endophytic, corticolous, or lichenicolous, and few are pathogenic species (Perera et al. 2023; Zhao et al. 2023; Zhao et al. 2025). Currently, this family comprises 50 accepted genera (Zhao et al. 2025), but whole-genome data remain limited. Presently, more than 30 genomes of this family have been sequenced and deposited in the NCBI database. Among these genomes, representative genera such as *Clonostachys*, *Emericellopsis*, *Geosmithia*, *Gliomastix*, *Hapsidospora,* and *Waltergamsia.* Nevertheless, the identification of BGCs within the *Bionectriaceae* family remains insufficiently explored. Some studies have identified BGCs in the sequenced genomes of this family. Research on *Clonostachys rosea* have reported a substantial array of genes encoding enzymes associated with secondary metabolites, including PKS, NRPS, and terpenes (Fatema et al. 2018; Wang et al. 2021). In *C. rosea* IK726, it was found that PKS gene expression is regulated by nutrient conditions and fungal interactions, with *pks29* contributing to antagonism against *Botrytis cinerea* (Fatema et al. 2018). Genomic analysis of the marine fungus *Emericellopsis atlantica* revealed its capability to produce a range of NRPS-derived peptides and polyketides, whereas in *Geosmithia*, a more limited BGC repertoire has been reported (Schuelke et al. 2017).

In this study, we sequenced and functionally annotated the genomes of two endophytic fungi of cacao to identify BGCs, in particular those potentially related to their antifungal activity, and to investigate how their biosynthetic gene cluster families (GCFs) compare with other members of the *Bionectriaceae* family. An exploratory analysis was conducted through genome mining to assess the biosynthetic potential of the two newly sequenced whole genomes, *Clonostachys rosea* TCF417 and *Waltergamsia zeylanica* TCF400, alongside the genomes of this fungal family available in public databases. Additionally, the similarity of BGCs was evaluated using the BiG-SCAPE engine (Navarro-Muñoz et al. 2020), which organizes the BGCs into gene cluster families (GCFs). Finally, extract fractionation and purification of verticillin D from *C. rosea* TCF417 was conducted followed by growth inhibition bioactivity evaluation of the fractions and purified molecule on two plant pathogens.

## Materials and methods

### Fungal culture and genomic DNA extraction

Two endophytic fungi strains previously isolated from young cacao pods, *W. zeylanica* TCF400 and *C. rosea* TCF417, were chosen based on their antagonistic activity towards cacao pathogen *Moniliophthora roreri* (Castillo et al. 2025). The source trees were found in La Magnita farm, Changuinola, Bocas del Toro Province, Panama, and the strains are currently deposited at the microbial culture collection of INDICASAT.

For genomic DNA extraction, the fungal strains were inoculated in 50 mL of Potato Dextrose Broth (PDB) (Difco 254920) and grown at 25°C, 125 rpm for 5-6 days in a shaking incubator. Mycelia were harvested by filtration using a sterile glass funnel with filter paper (Whatman N°1, Whatman International Ltd., Maidstone, England). The biomass was lyophilized (Labconco, Kansas, USA) and stored at -80 °C. Genomic DNA was extracted following the protocol instructions for DNA Purification from tissue using Gentra Puregene Yeast/Bact. Kit (QIAGEN, Hilden, Germany). DNA quality and concentration were evaluated using NanoDrop 2000 (Thermo Fisher Scientific, Waltham, MA, USA) and a Qubit fluorometer (Thermo Fisher Scientific, Waltham, MA, USA). In addition, DNA conditions were determined using 1% agarose gel electrophoresis.

### Library Preparation and Sequencing

Genomic DNA Library of *W. zeylanica* TCF400 and *C. rosea* TCF417 were prepared using the Nanopore Genomic DNA by ligation kit, SQK-LSK109 and SQK-LSK114 respectively, from Oxford Nanopore Technologies (ONT, Oxford, UK) following the manufacturer’s instructions. Library sequencing was performed on an Oxford Nanopore MinION Mk1C sequencer using a FLO-MIN-106 R9.4.1 and FLO-MIN-114 R10 flow cell (ONT, Oxford, UK) at the Genomics Laboratory of INDICASAT AIP (Panama). MinKNOW (version 21.02.2, 23.07.12 ONT) was used to control the run using the 72-h sequencing run protocol. High accuracy basecalling was performed in real-time using Guppy (version 4.3.4 and 7.1.4). Nanopore quality control was achieved using MinIONQC v1.4.1 with a threshold quality (Q) value > 7 (Lanfear et al. 2019). Nanopore reads were trimmed using Porechop v0.2.4 (Wick 2018).

Illumina genome sequencing of *W. zeylanica* TCF400 was performed by Macrogen Inc. (Seoul, Korea). A 350 bp DNA library (Illumina TruSeq DNA PCR-Free) followed by a 150 bp paired-end sequencing using the Illumina Novaseq sequencing platform (Illumina, San Diego, CA, USA). *Clonostachys rosea* TCF417 was sequenced with Illumina technology at the Genomics Laboratory of INDICASAT AIP. More specifically, a DNA library of 500 bp (Illumina TruSeq DNA PCR-Free) was employed for a 250 bp paired-end sequencing in the Illumina MiSeq platform (Illumina, San Diego, CA, USA).

### Genome Assembly and Annotation

Genome assembly was performed *de novo* using ONT long reads with Canu 2.2 assembler (Koren et al. 2017). The resulting genome assembly was polished by three iterations on Racon 1.4.22 (Vaser et al. 2017) and one iteration using Medaka 1.7.1 (nanoporetech/medaka 2024). Illumina paired-end reads were quality trimmed using Trimmomatic 0.39 (Bolger et al. 2014). The high-quality trimmed Illumina reads were aligned to the Canu genome assembly using BWA 0.7.17-r1188 (Li and Durbin 2009) and filtered using samtools 1.11 (Li et al. 2009). The genome assembly was then corrected with high-quality Illumina alignments using three iterations of Pilon 1.24 (Walker et al. 2014). Genome assembly was evaluated using QUAST 5.0.2 (Mikheenko et al. 2018). For annotation of the draft genome assemblies, we used Funannotate v1.8.15 (Palmer and Stajich 2020) available on the Galaxy web platform (The Galaxy Community 2024). Protein and transcript data were obtained from the Joint Genome Institute database, from *Acremonium chrysogenum* ATCC 115501 (Terfehr et al. 2014), currently known as *Hapsidospora chrysogena* (*Bionectriaceae*) (Hou et al. 2023). As Funannotate uses Benchmarking Universal Single-Copy Orthologs (BUSCO) for the initial training of *ab initio* predictors, we selected sordariomycetes_odb10 and *Fusarium graminearum* as the seed species for BUSCO searches. Functional annotations from eggNog Mapper (v2.1.8) and InterproScan (v5.59-91.0) were integrated with the command funannotate annotate. Subsequently, annotated genomes were evaluated using BUSCO (Manni et al. 2021).

### BGC identification and gene cluster family networks

To gain insight into the biosynthetic potential of *W. zeylanica* TCF400 and *C. rosea* TCF417, secondary metabolite gene clusters were identified using the fungal version of antiSMASH 7.1.0 in strict mode to minimize the detection of fragmented BGCs (regions) resulting from differences in genome assemblies, with CASSIS implemented for cluster boundary prediction. In antiSMASH output, a “region” refers to a predicted biosynthetic gene cluster (BGC) (Blin et al. 2023). Additionally, a total of 30 publicly available fungal genomes from the *Bionectriaceae* were included. These were retrieved from NCBI as of August 2024, representing six different genera *Clonostachys*, *Emericellopsis*, *Geosmithia*, *Gliomastix*, *Hapsidospora* and *Waltergamsia* (Supplementary Table 1). To compare gene sequences and BGCs identified via AntiSMASH and determine homologous and widely distributed BGCs whole dataset, we used the Biosynthetic Gene Similarity Clustering and Prospecting Engine (BiG-SCAPE) (Draisma et al. 2026). BiG-SCAPE v2.0.0 was run in cluster mode with Pfam v37.2 (Mistry et al. 2021), and BGC network was generated by applying gene cluster family (GCF) cutoffs of 0.3, 0.4, 0.5, and 0.6. To prevent overestimation of GCF, we chose the network with a cutoff of 0.5. Other parameters used: --classify class, --alignment-mode auto, singletons were not included. The resulting similarity networks were visualized with Cytoscape v3.10.3 (Otasek et al. 2019). Using the CAGECAT platform (Belt 2023), homology analysis and synteny visualization of multiple BGCs were conducted utilizing Clinker and Clustermap.js (Gilchrist and Chooi 2021), respectively. The analysis focused on representative GCFs from the most prevalent BGC categories (PKS, NRPS, terpenes and hybrid NRPS-T1PKS) specifically those that showed similarity to entries in the MIBiG database.

### Fractionation of the extract of Clonostachys rosea TCF417 and isolation of Verticillin D

The fungal strain *C. rosea* TCF417 was cultivated separately on 200 plates of malt extract agar and incubated for 31 days at room temperature (25 ± 2°C). The fungal culture was sonicated and macerated twice with 1200 mL of ACS-grade ethyl acetate (EtOAc) for 24 h with constant agitation (Boya P. et al. 2024). The organic extract was filtered through a 1.0 μm fiberglass filter and dried under reduced pressure to yield 1.29 g of the extract. The crude EtOAc extract of *C. rosea* TCF417 was fractionated using open-column chromatography with silica gel as the stationary phase. Elution was performed using a gradient system of 20, 40, 60, 80, and 100% EtOAc in hexane, followed by 20 and 40% methanol in EtOAc, resulting in seven fractions (F1–F4). The fraction F4 (80% EtOAc in hexane) was dissolved in methanol (1 mg/20 µL) and purified via high-performance liquid chromatography (HPLC) using a semipreparative C18 reverse-phase column (Synergi Hydro-RP 10 mm × 250 mm), which was eluted with a gradient system of 70–80% methanol in water for 25 min, followed by isocratic elution with 100% methanol for 35 min, resulting in the purification of verticillin D.

### Antifungal Activity

The *in vitro* antifungal activity of fractions (F1-F4) was evaluated using the poisoned food bioassay method (Rodriguez et al. 2024). Briefly, samples were dissolved in dimethyl sulfoxide (DMSO) and added to potato dextrose agar (PDA) medium previously cooled to 40 °C to obtain a final concentration of DMSO in PDA as 0.5% V/V. Then, 5 mL of the poisoned medium was poured to 6 cm diameter Petri dishes. A plug of mycelium from actively growing culture of the fungal pathogens *Moniliophthora roreri* LCM1133 and *Mycena citricolor* MC01 in PDA were taken with a 5 mm hole puncher and inoculated at the center of the plates. These isolates were obtained from the microbial culture collection of INDICASAT. The treatments were incubated at 25 °C (Li et al. 2019) using five replicate per fraction for *Moniliophthora roreri* and three replicates for *Mycena citricolor*. Bioactivity was determined as growth inhibition percentages. Mycelial growth of fungal pathogens was measured using the cross-bracket method, and growth inhibition percentages (GI%) were calculated as GI %: (1 − T/C) × 100, where C corresponds to the radial growth of the pathogen in the negative control and T is equal to the radial growth of the pathogen under treatment (Rashid et al. 2018). Mancozeb (MNZ) was dissolved in DMSO, added to a final concentration of 50 µg/mL, and used as a positive control for the growth inhibition assay. As a negative control, we employed DMSO at 0.5% V/V. Column fractions (CF1–CF4) were dissolved in DMSO (final concentration 0.5% V/V) and evaluated at 250 µg/mL (Balouiri et al. 2016).

The *in vitro* antifungal activity of Verticillin D against *Moniliophthora roreri* and *Mycena citricolor* was evaluated using the agar disk diffusion method (Bonev et al. 2008). Verticillin D was dissolved in DMSO at a concentration of 100 µg/10 µL and 10 µL was applied to a 5 mm of diameter cellulose disk placed 1 cm from a plug of mycelium of the fungal pathogen. Five replicates were used per treatment for each fungal pathogen. Growth inhibition was assessed at 3, 5, 7, and 13 days.

### Statistical Analysis

Growth inhibition percentage (GI%) was calculated from measurements taken on days 3, 5, 7, 10, and 13. Statistical comparisons were performed separately for *Moniliophthora roreri* and *Mycena citricolor* at days 7 and 13, corresponding to the peak and final evaluation points, respectively. For the fraction assay, F1–F4 were compared with MNZ and DMSO, while verticillin D was compared with the same controls. One-way ANOVA was used for treatment comparisons, followed by Tukey’s honestly significant difference (HSD) test when the ANOVA was significant (p < 0.05). Measurements from days 3 and 5 were used to monitor the progression of growth inhibition but were not included in the statistical analysis. All analyses and visualizations were performed in R v4.2.0 using dplyr, agricolae, ggplot2, and cowplot. Compact letter displays were generated using agricolae to indicate significant differences among treatments.

## Results

### General features of sequenced genomes

Two species of *Bionectriaceae* were sequenced in this study: *C. rosea* TCF417 and *W. zeylanica* TCF400, using ONT and Illumina sequencing technologies. The genome assembly sizes were 52.9 Mb and 31.0 Mb, for *C. rosea* TCF417 and *W. zeylanica* TCF400 respectively (Table 1). The genomes of the strains were similar in size to previously sequenced genomes of *Bionectriaceae* publicly available in the NCBI database, which had an average genome size of 49.11 Mb (Supplementary Table S1). BUSCO estimate of genome completeness using the fungi_odb10 database resulted in 97.1% for the genome assembly of *C. rosea* TCF417 and 99.0% for *W. zeylanica* TCF400 (Table 1, Supplementary Table 2), these results showed high level of completeness for both assemblies.

**Table 1.** Summary of genome assembly and annotation* of *Clonostachys rosea* TCF417 and *Waltergamsia zeylanica* TCF400.

| **Genome statistics** | **Organism** |  |
|---|---|---|
|  | ***Clonostachys rosea* TCF417** | ***Waltergamsia zeylanica* TCF400** |
|---|---|---|
| Genome size (Mbp) | 52.9 | 31.0 |
| GC% | 50.30 | 53.04 |
| N50 | 190,966 | 3,522,015 |
| scaffolds | 469 | 24 |
| BUSCO completeness (%) | 97.10% | 99.0% |
| Number of genes | 17,030 | 10,454 |
| mRNA | 16,810 | 10,288 |
| tRNA | 220 | 166 |
| average gene length | 1,619 | 1,589 |
| GO terms | 10,680 | 7,161 |
| InterProScan | 13,325 | 8,493 |
| eggNOG | 15,763 | 9,890 |
| Pfam | 11,615 | 7,501 |
| CAZyme | 797 | 451 |
| MEROPS | 635 | 397 |
| BUSCO | 3,735 | 3,751 |
| Accession number | JBICNQ000000000 | JBICNP000000000 |

\*Funannotate uses predicted protein models to perform functional annotation through several tools, including InterProScan, and Eggnog-mapper. It also conducts alignment-based annotation using reference databases such as Pfam 35.0, Uniprot 2023\_01, MEROPS 12.0, dbCAN 11.0, MIBiG 1.4, and interPro 93.3. These databases help identify Pfam domains, CAZymes, secreted proteins, peptidases (MEROPS), and BUSCO groups.

We identified a total of 17,030 genes in *C. rosea* TCF417 and 10,454 genes in *W. zeylanica* TCF400, with average lengths of 1,619 bp and 1,589 bp, respectively (Table 1). The number of mRNA features was 16,810 for *C. rosea* TCF417 and 10,288 for *W. zeylanica* TCF400, with tRNA counts of 220 and 166, respectively. Functional annotation indicated 10,680 and 7,161 Gene Ontology (GO) terms, 13,325 and 8,493 InterProScan matches, and 15,763 and 9,890 eggNOG annotations for *C. rosea* TCF417 and *W. zeylanica* TCF400, respectively. Furthermore, 11,615 and 7,501 genes were assigned to the Pfam domains for *C. rosea* and *W. zeylanica*. In terms of specialized gene families, *C. rosea* TCF417 had a higher number of carbohydrate-active enzymes (CAZymes; 797) and proteases (MEROPS entries; 635) compared to *W. zeylanica* TCF400 (451 CAZymes and 397 MEROPS). Information regarding the genome assembly and annotation is summarized in Table 1 and Supplementary Table 3.

### Biosynthetic gene cluster distribution

Biosynthetic genes clusters (BGCs) were predicted in all 32 genomes analyzed in this study using the fungal version of antiSMASH. This analysis identified a total of 1,523 regions that contain BGCs (Figure 1 and supplementary Table 4). The majority of these BGCs belong to the type I polyketide synthases (T1PKSs) category (591 clusters), followed by non-ribosomal peptide synthases (NRPSs) (331 clusters), terpenes (202 clusters) and hybrid NRPS-T1PKS clusters (150 clusters).

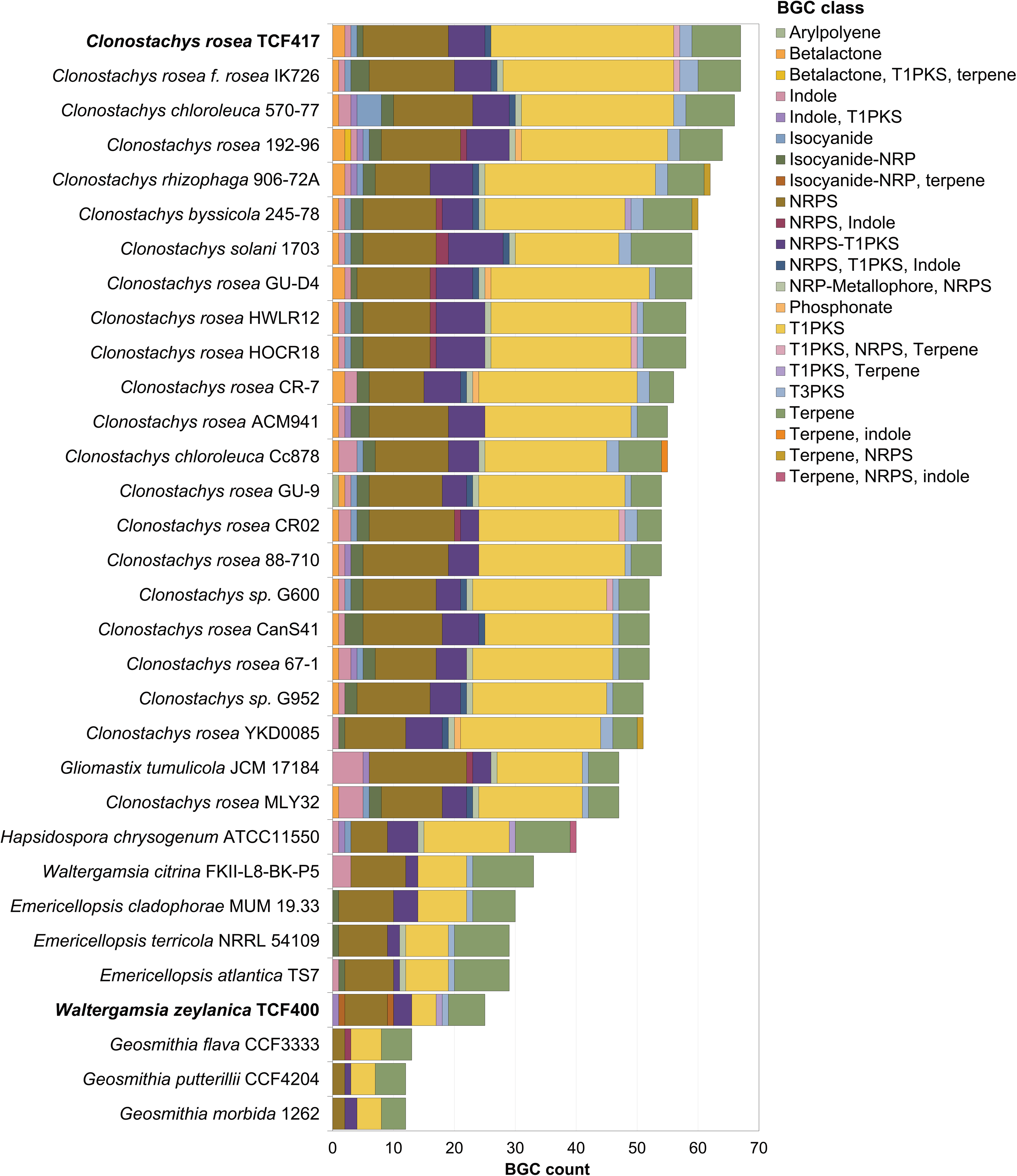

Among the analyzed genomes, *C. rosea* TCF417 and *C. rosea* IK726 had the highest number of predicted BGCs, with 67 clusters each (Figure 1). Specifically, we found that *C. rosea* TCF417 showed the highest number of T1PKS clusters (30 BGCs), surpassing all other members of the genus *Clonostachys* and other taxa within the *Bionectriaceae* family (Figure 1 and supplementary Table 4). In the NRPS category, *Gliomastix tumulicola* JCM 17184 had the greatest number of BGCs (16 clusters), while species of *Geosmithia* had the fewest, with two NRPS BGC per each genome (Supplementary Table 4). For terpene biosynthetic gene clusters, it was found that *C. solani* 1703 had the highest number (10 BGCs), whereas *C. rosea* CR02, CR-7 and YKD0085, along with *W. zeylanica* TCF400 showed the lowest number of terpene clusters, with four BGCs per each genome. The NRPS-T1PKS hybrid category was the fourth most abundant across the *Bionectriaceae*. *Clonostachys rosea* HOCR18 and HWLR12 each contained eight BGCs of this type, while *E. atlantica* TS7 and *G. putterillii* CCF4204 each had one NRPS-T1PKS hybrid cluster.

### Biosynthetic gene cluster similarity networks

To characterize the homologous gene clusters in the *Bionectriaceae* family, gene networks for each BGC category were generated using the BiG-SCAPE program. Within each network, similar BGCs were grouped into gene cluster families (GCFs). A total of 209 GCFs were predicted for *Bionectriaceae*, including the following categories: betalactone, indole, isocyanide, isocyanide-NRP, NRP-metallopore-NRPS, NRPS, NRPS-indole, NRPS-T1PKS, NRPS-T1PKS-indole, NRPS-T1PKS-terpene, phosphonate, T1PKS, T1PKS-indole, T3PKS, and terpene. A total of 52 GCFs contained previously characterized BGCs from the MIBiG database, but only 11 GCF clearly included BGCs identified in cacao endophytes genomes and MIBiG reference BGCs. For the most outstanding BGC categories (T1PKS, NRPS and NRPS-T1PKS and terpene), we found a total of 98 GCFs for T1PKS which includes 34 GCFs with characterized BGCs, most these GCFs contained only characterized BGCs from MIBiG database, without representatives from our genomes (Figure 2, supplementary Table 5). For NRPS, there were 39 GCFs with 4 of them having characterized BGCs (Figure 3, supplementary Table 5). Meanwhile, both NRPS-T1PKS and terpene had 22 GCFs each, with 7 and 4 characterized BGCs, respectively (Figure 4, Table S5). A summary of biosynthetic gene cluster families (GFC) identified in the dataset using BiG-SCAPE is showed in supplementary tables 5 and 6.

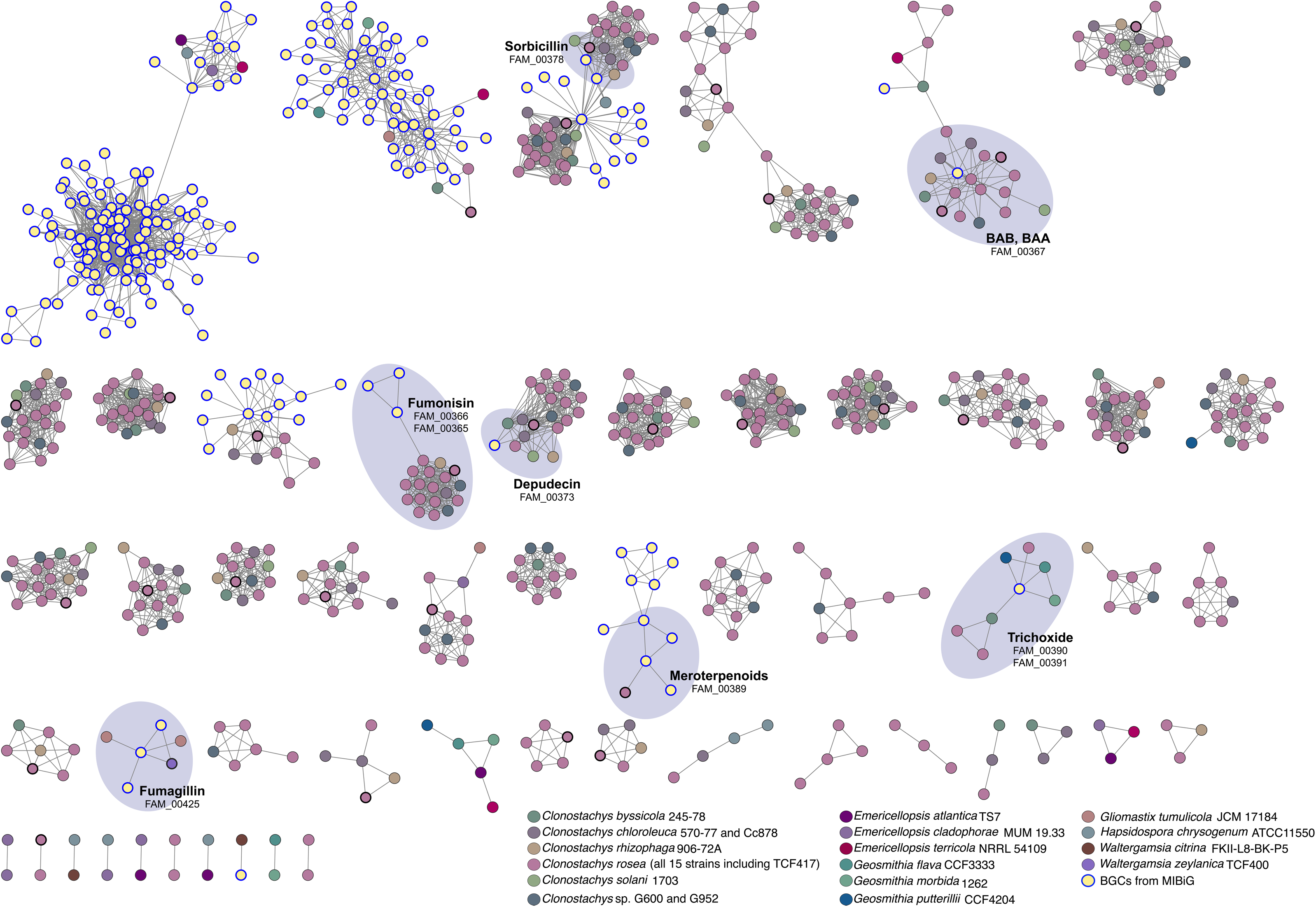

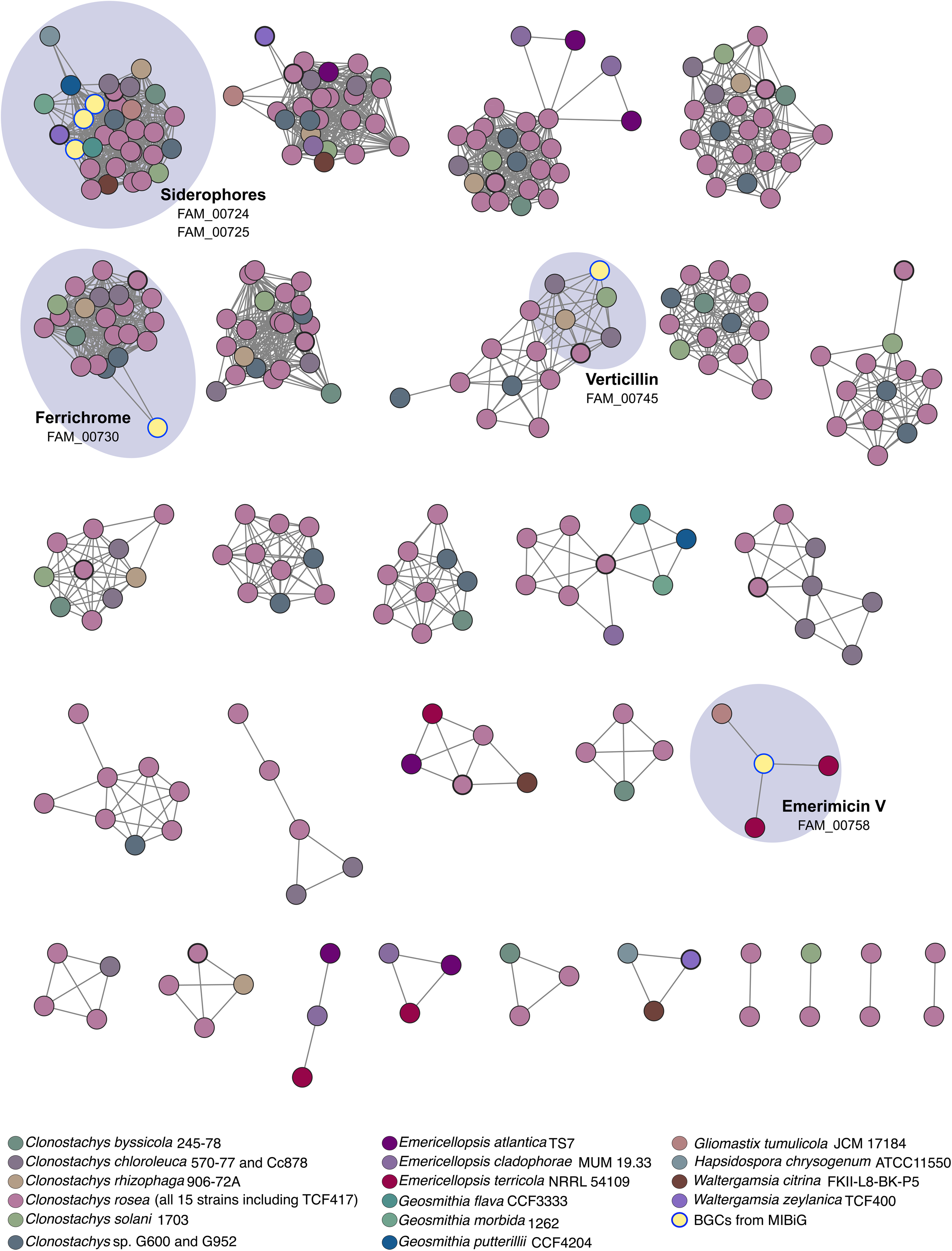

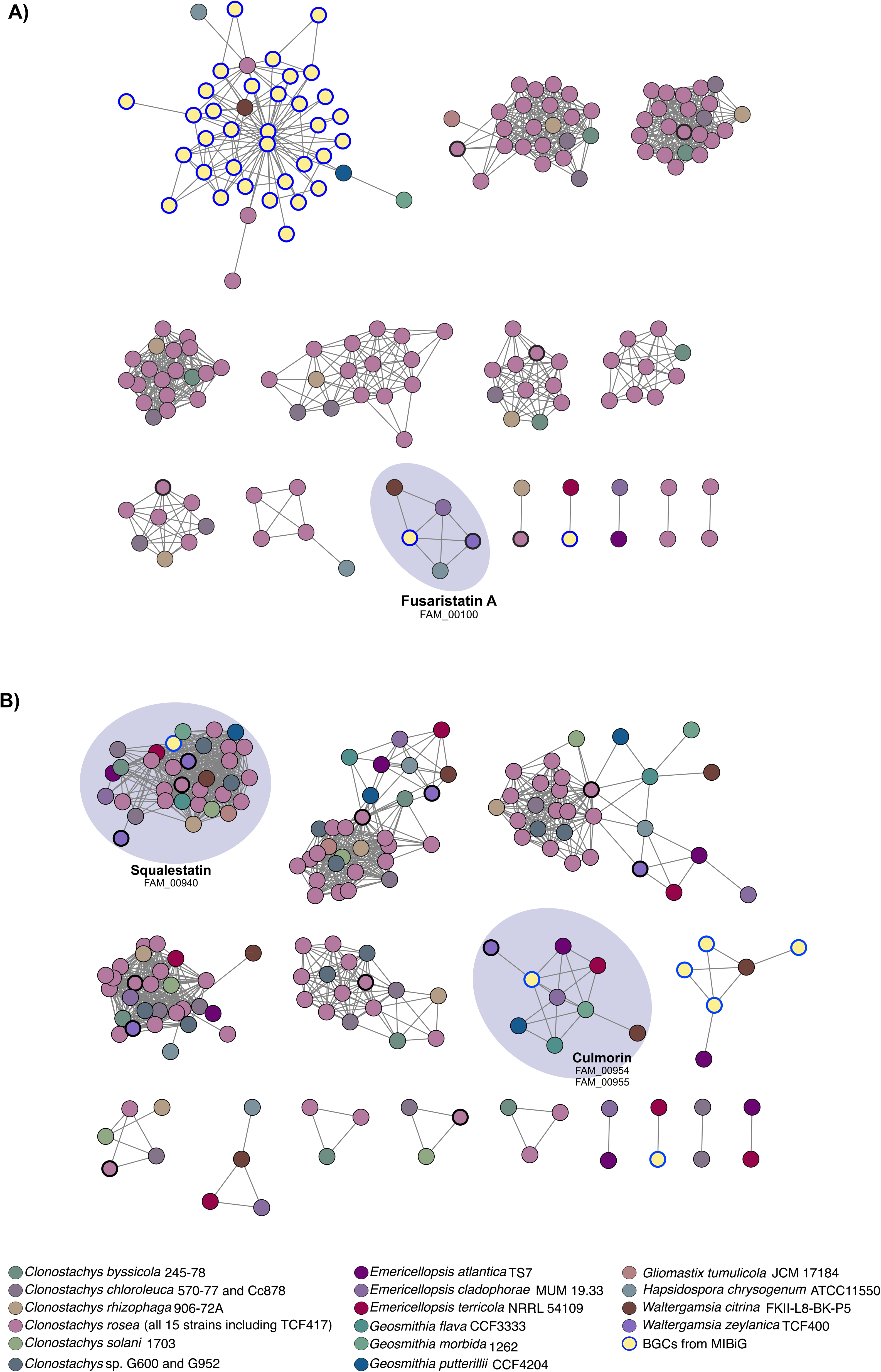

### Type 1 PKS gene cluster family network and synteny analysis of sorbicillinoids

In the T1PKS dataset (Figure 2), most of GCFs (98) could not be linked to known products, indicating a large proportion of uncharacterized biosynthetic diversity in *Bionectriaceae*. Only 5 GCFs included BGCs from cacao endophytes genomes. Several GCFs were associated with known metabolite families. For example, FAM_00365 grouped BGCs from *C. rosea* TCF417 with other *Clonotachys* species, and was closely related to FAM_00366, which includes fumonisin clusters (BGC0000062, BGC0000063, BGC0002502). However, the identified BGCs from FAM_00365 mainly share PKS-related domains with these reference clusters and lack the complete gene set required for fumonisin biosynthesis, suggesting only partial similarity. In addition, two BGCs from *C. rosea* TCF417 (regions 122.1 and 267.1) were assigned to FAM_00367, corresponding to BGC0002240, which encode the compounds (2*E*,4*E*,6*E*,10*E*)-9-hydroxydodeca-2,4,6,10-tetraenoic acid (BAB), and (2*Z*,4*E*,6*E*,10*E*)-9-hydroxydodeca-2,4,6,10-tetraenoic acid (BAA). This GCF also includes homologous regions from other *Clonostachys* species, such as: *C. rosea*, *C. rhizophaga*, *C. chloroleuca,* and *C. solani.* Other notable association included meroterpenoid-related clusters in FAM_00389 (e.g.: paraherquonin, novofumigatonin, emeridone F, and anditomin), which includes the region 405.1 from *C. rosea* TCF417 genome. Other GCFs association with characterized clusters included, FAM_00425 for fumagillin-related clusters in which regions from *W. zeylanica* TCF400 and *G. tumilicola* JCM 17184 were assigned. Also, trichoxide-associated clusters were identified in FAM_00390 and FAM_00391 across multiple genera. A conserved GCF (FAM_00373) linked to depudecin was detected across several *Clonostachys* species. Similarly, sorbicillin-associated clusters (BGC0001404.3, BGC0002190.2) in FAM_00378, included a BGC from *C. rosea* TCF417 and related *Clonostachys* species. Synteny analysis of this GCF revealed a conserved backbone genes across strains with variations mainly in gene organization and accessory gene content (Figure 5).

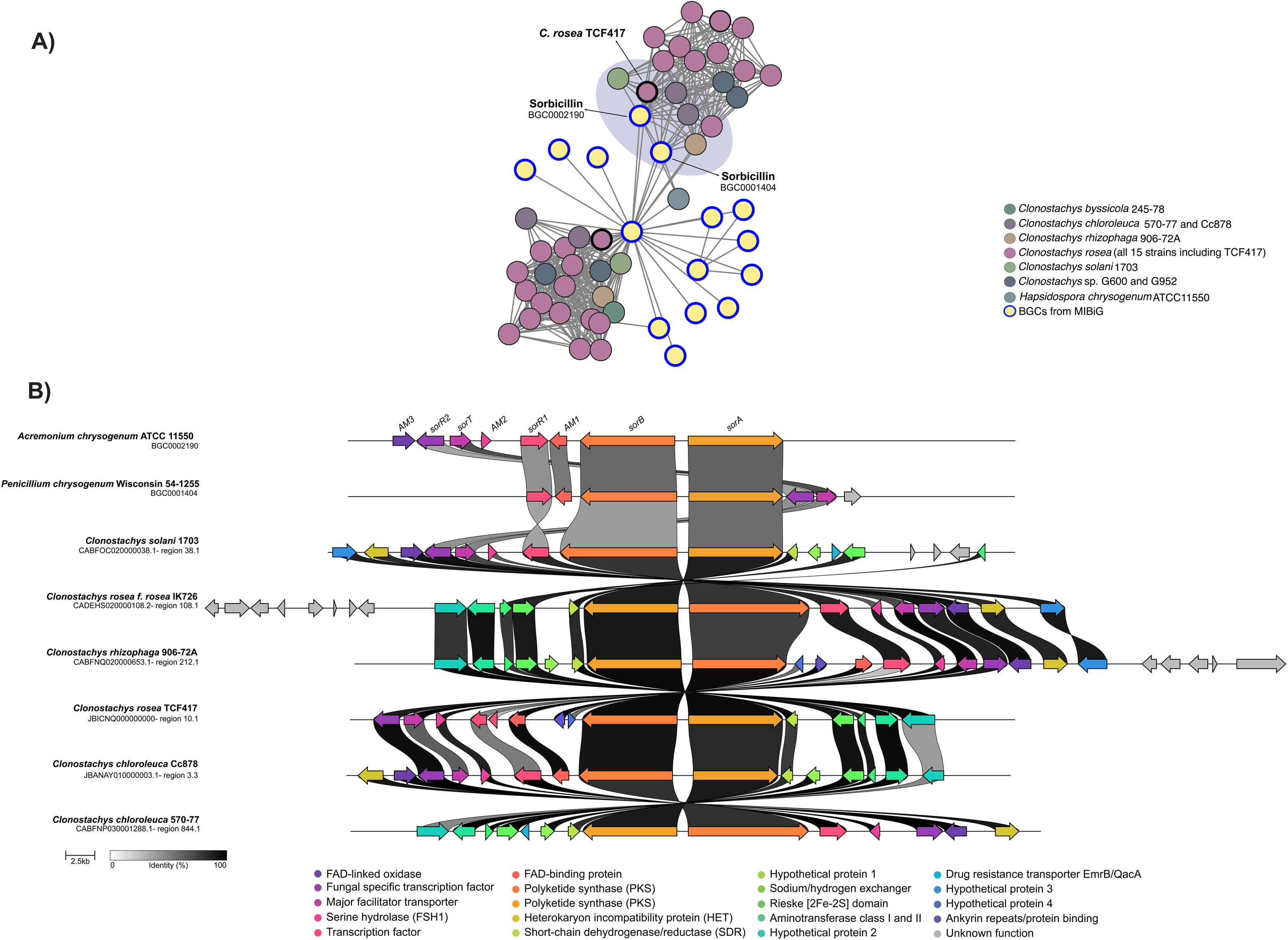

### NRPS gene cluster family network and synteny analysis of verticillin

Similar to the T1PKS dataset, most NRPS GCFs could not be assigned to a specific product or product family (Figure 3). However, some conserved and relevant clusters were identified. FAM_00724, FAM_00725, and FAM_00730 were associated with siderophore biosynthesis, including ferrichrome (BGC0000900, BGC000090), dimethylcoprogen (BGC0001249), and metachelin C (BGC0002710). These clusters were widely distributed across *Bionectriaceae* including *C. rosea* TCF417 and *W. zeylanica* TCF400 and multiple regions from different genera such as: *Clonostachys*, *Geosmithia*, *Hapsidospora*, and *Gliomastix*.

The verticillin BGCs (FAM_00745) were also identified across *Clonostachys* species, including *C. rosea* TCF417 (Figure 6A). Synteny analysis showed a mostly conserved gene organization and high sequence similarity compared to the reference BGC for verticillin (BGC0001820). However, the cluster identified in *C. rosea* TCF417 contained fewer genes than those in closely related strains (Figure 6B). Additionally, a BGC for emerimicin V (BGC0002555) was identified in *E. terricola* NRRL54109 and *G. tumulicola* JCM17184, indicating a broader distribution of NRPS-derived metabolites within the family.

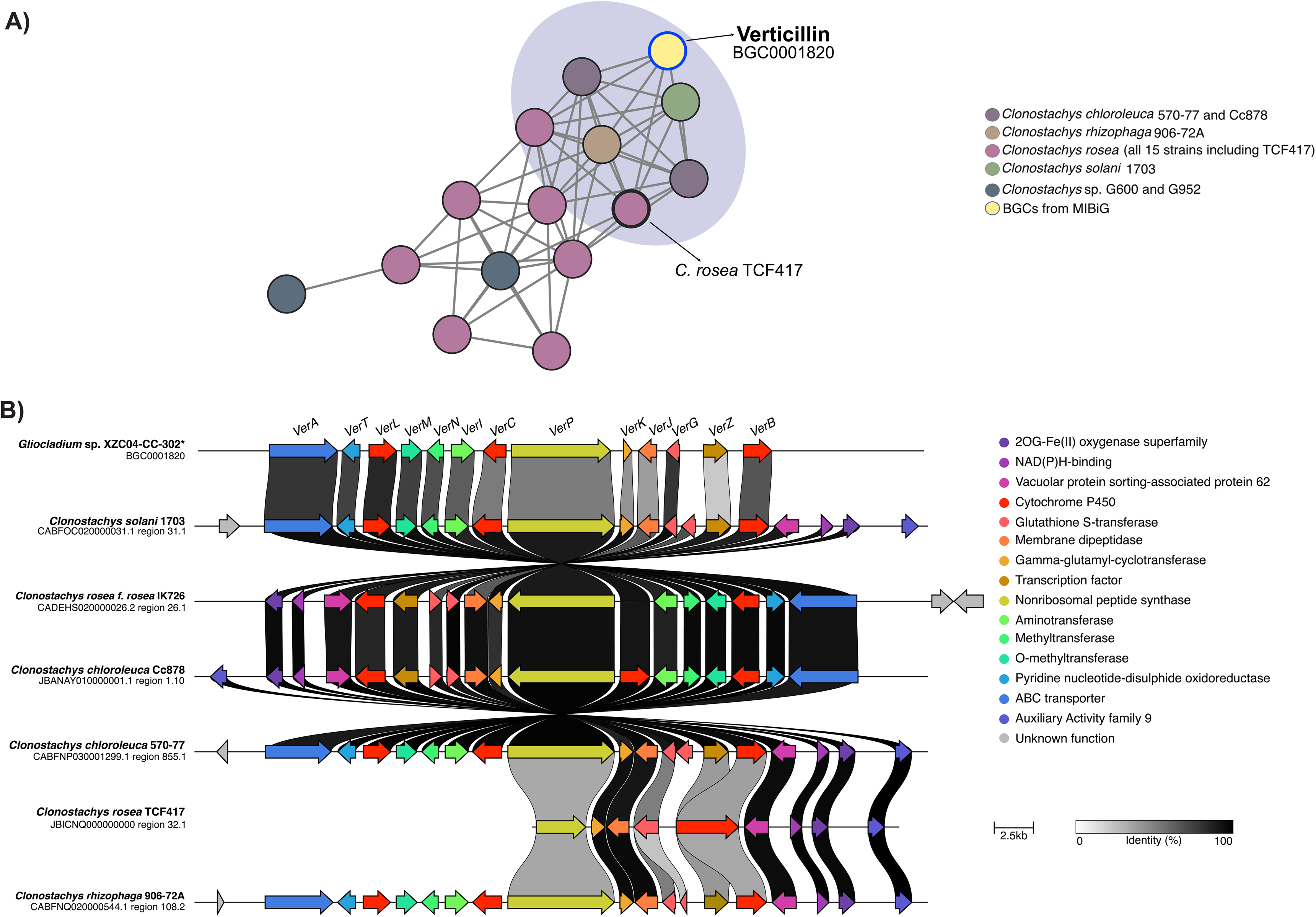

### NRPS-T1PKS gene cluster family network and synteny analysis of fusaristatin A clusters

In the hybrid NRPS-T1PKS, a limited number of characterized BGCs were associated with GCFs (Figure 4A). Among these, fusaristatin A (BGC0002186; FAM_00100), a lipopetide of hybrid origin, was identified in *W. zeylanica* TCF400, as well as in *E. cladophorae* MUM19.33, *H. chrysogenum* ATCC11550, and *W. citrina* FKII-L8-BK-P5 (Figure 4A and Figure 7A). Synteny analysis showed that these clusters share conserved core genes (NRPS7 and PKS6), along with accessory genes such as cytochrome P450 monooxygenase, aminotransferase, and a stress-responsive protein. Notably, genes of unknown function exhibited low conservation across strains (Figure 7B).

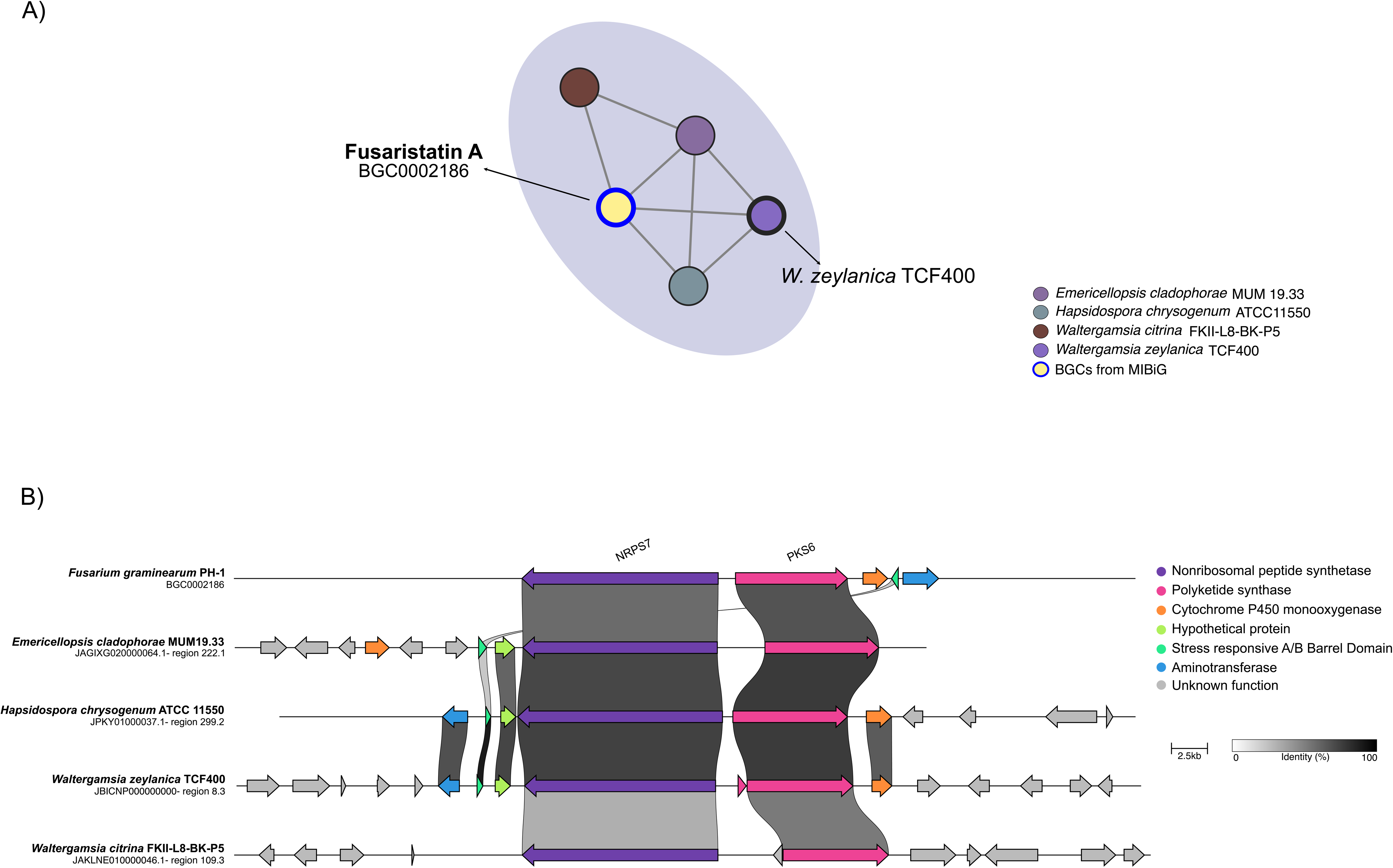

### Terpene gene cluster family networks and synteny analysis of squalestatin S1

In the terpene dataset, the culmorin BGC (BGC0002176) was identified in FAM_00954 and FAM_00955, including *W. zeylanica* TCF400 and species from *Emericellopsis*, *Geosmithia*, and *Waltergamsia* (Figure 4B).

The squalestatin S1 cluster (BGC0001839) was widely distributed across *Bionectriaceae*, with multiple strains from different genera grouped within the same GCF (Figure 4B), such as *Clonostachys* (including *C. rosea* TCF417), *Geosmithia*, *Gliomastix*, *Emericellopsis*, and *Waltergamsia*. Notably, two BGCs from *W. zeylanica* TCF400 were included in this cluster. Synteny analysis of the squalestatin S1 GCF revealed a conserved gene content and organization across strains. In addition to the core squalene/phytoene synthase gene, conserved genes included DnaJ domain-containing protein, hypothetical protein 1 and peptidase S8 domain gene (Supplementary Figure 1).

### Antifungal activity of fractions and verticillin D from *Clonostachys rosea* TCF417

The fractions F1-F4 and the compound verticillin D obtained from *C. rosea* TCF417 were evaluated against *Moniliophthora roreri* LCM1133 and *Mycena citricolor* MC01. Among the tested fractions, F4 and F1 exhibited the highest bioactivity on day 7 against *Mycena citricolor*, achieving 95.2% and 86.7% mycelial growth inhibition (GI%), respectively, which was statistically comparable to the commercial fungicide Mancozeb (99.0%, Tukey group a) (Figure 8A, supplementary Tables 7 and 8). In contrast, against *Moniliophthora roreri*, mancozeb showed the highest inhibition (91.8%; group a), followed by F4 (45.3%; group b). On day 13, F4 retained substantial activity against *Mycena citricolor* (86.4%, group b), whereas the activity of F1–F3 progressively decreased (Supplementary material: Figure S2, Tables S7 and S8).

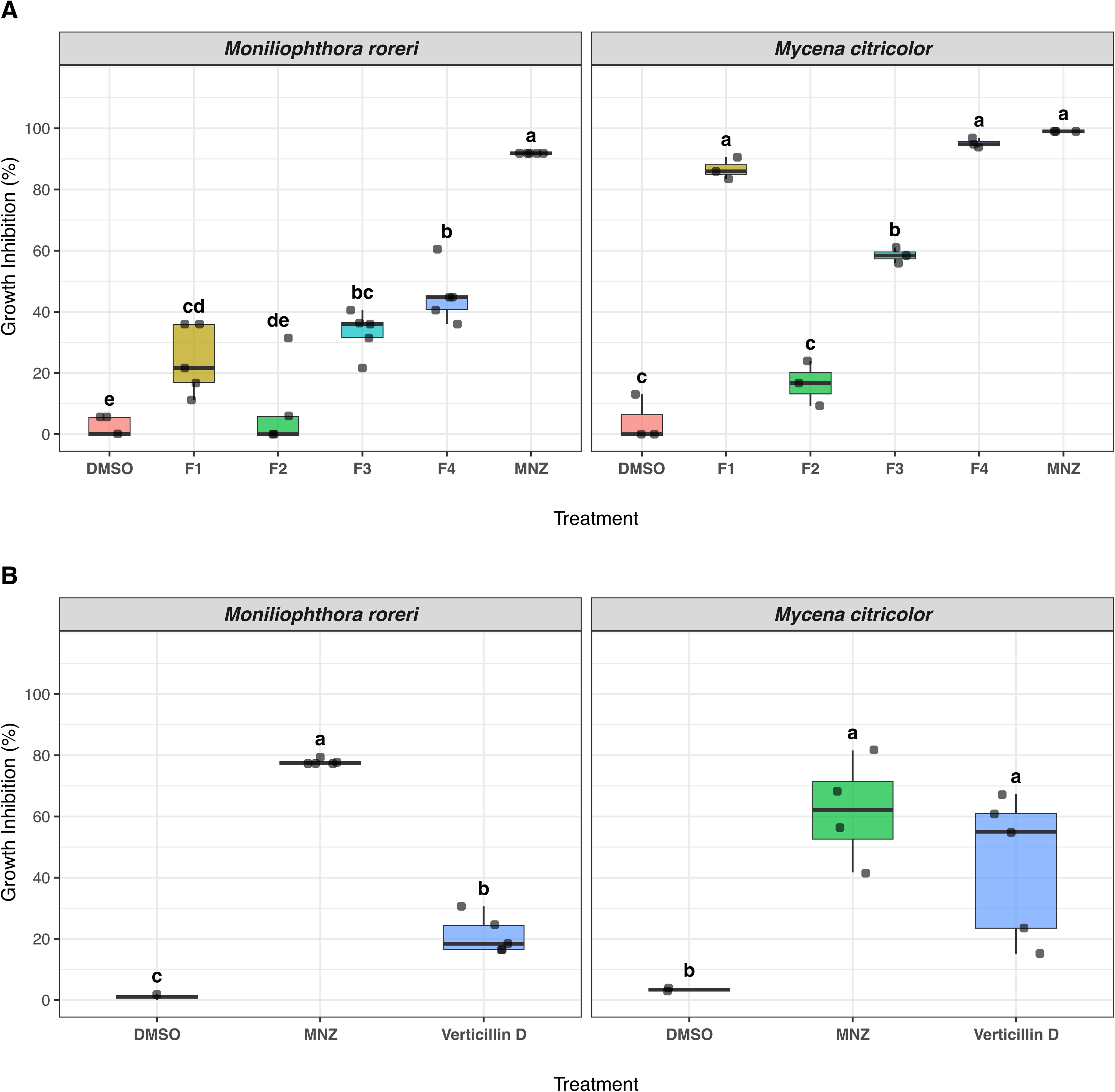

The agar disk diffusion assay revealed a species-specific response to verticillin D, previously isolated from F1 (Boya P. et al. 2024). Against *Moniliophthora roreri*, verticillin D exhibited moderate and relatively stable growth inhibition, ranging from 21% to 28%, with the highest inhibition observed on day 10 (Supplementary Table 9). On day 7, verticillin D showed significantly greater inhibition than the negative control (p < 0.05; Figure 8B; Supplementary Tables 7 and 10) and maintained approximately 21% inhibition on day 13. On the other hand, verticillin D showed remarkable bioactivity against *Mycena citricolor,* reaching its maximum inhibitory activity on day 7 (44 ± 24%, supplementary Table 9). Post-hoc mean comparisons using Tukey’s HSD test showed that the bioactivity of verticillin D on day 7 was statistically comparable to mancozeb against *Mycena citricolor* (Tukey group a; Figure 8B, supplementary Table 10). By day 13, the three treatments were assigned to distinct statistical groups (Supplementary material: Figure S3B, Table S10).

Together, these findings demonstrate that, fractions and verticillin D exhibited measurable antifungal activity against the two phytopathogens. Although their overall inhibitory activity was mild to moderate compared to the commercial fungicide mancozeb, their activity, particularly against *Mycena citricolor*, highlights their potential as source of antifungal compounds against these fungal pathogens.

## Discussion

The genome mining of available genomes of *Bionectriaceae* conducted here, including the newly sequenced genomes of *C. rosea* TCF417 and *W. zeylanica* TCF400, revealed a high diversity of biosynthetic gene clusters, highlighting the substantial metabolic potential of this fungal family. A total of 1,523 BGCs identified, grouped into 209 GCF, 52 of which were linked to previously characterized BGC (∼24%). However, many of these pathways remain hidden due to a lack of corresponding entries in reference databases such as MIBiG. This large fraction of unassigned GCF (∼75%) indicates that much of the biosynthetic capacity of *Bionectriaceae* remains unexplored, suggesting a rich source of secondary metabolites. Previous research has indicated the need to utilize genome data to understand the biosynthetic potential of *Bionectriaceae* (Zhao et al. 2025). This study provides the first insight into the biosynthetic gene clusters harbored in publicly available genomes of *Bionectriaceae* including two endophytic fungi from cacao here sequenced and BGC similarity network analyzed.

The predominance of PKS, NRPS, hybrids NRPS-T1PKS and terpenes clusters is consistent with patterns observed across Fungi (Robey et al. 2021; Zhang et al. 2024), where T1PKS category is the most abundant, indicating that a polyketides structure varies greatly between species, although some species are taxonomically closely related (Zhang et al. 2024). In particular, the dominance of T1PKS clusters aligns *Bionectriaceae* with families such as *Hypoxylaceae*, where a total of 783 BGCs were identified, grouped into 375 GCFs based on 14 genomes analyzed, and most of the clusters were identified as PKS (Kuhnert et al. 2021). Whereas other families of fungi like *Pleosporaceae* show a higher prevalence of NRPS clusters with a total of 6,323 BGCs reported grouped into 548 GCFs across 187 genomes (Kim and Dettman 2025). These differences highlight how both taxonomy and sampling depth shape the observed biosynthetic landscape. In large-scale genome mining studies, the potential chemodiversity encoded in fungal genomes has been estimated at over 1.4 million of secondary metabolites (SMs), considering that a single BGC can biosynthesize multiple compounds (Robey et al. 2021; Zhang et al. 2024; Riedling and Rokas 2025). Although these studies have identified some taxonomic groups within the subphylum Pezizomicotina with a higher number of BGCs and/or SMs, their datasets include small numbers of species from the family *Bionectriaceae* (Robey et al. 2021).

Among the GCFs linked to known metabolites, sorbicillinoids represent a well-characterized PKS-derived group that illustrates the functional diversity encoded in these genomes. Sorbicillinoids are natural products derived from diverse fungi, including members of the orders Hypocreales and Eurotiales, such as *Trichoderma* and *Penicillium*, respectively (Chen and Chu 2019; M. Milzarek and M. Gulder 2025). A broad range of bioactivities have been reported for this group of molecules, including antibacterial, antifungal, cytotoxic, and anti-inflammatory activities (M. Milzarek and M. Gulder 2025). The biosynthetic gene cluster for sorbicillinoids has been described in *Penicillium chrysogenum* E01-10/3 and *P. rubens* Wisconsin 54-1255 (Fahad et al. 2013; Salo et al. 2016), *Trichoderma reesei* (Derntl et al. 2017), and *Acremonium chrysogenum* ATCC 11550 (Chen and Chu 2019). The sorbicillinoid gene cluster harbors several genes, including two polyketide genes (SorbA and SorB), which are responsible for their core structure (Fahad et al. 2013; Salo et al. 2016; Chen and Chu 2019). According to our results, sorbicillinoids clusters showed conserved core biosynthetic genes alongside variations in accessory genes. These differences may indicate potential structural diversification in the molecules produced by each strain. In a previous study with *C. rosea* TCF417, three metabolites from this molecular family were reported: bislongiquinolide, trichodimerol and dihydrotrichodimerol (Boya P. et al. 2024). These compounds were dereplicated using the feature based molecular networking approach; therefore, their activity was not evaluated (Boya P. et al. 2024). Some sorbicillinoids have been evaluated against plant pathogens for example, compounds from *Trichoderma* spp. showed activity against cacao pathogens *M. perniciosa* and *M. roreri* (Chóez-Guaranda et al. 2023). Similarly, antifungal extracts from *C. rosea* CMAA1284 were effective against gray mold on cherry tomatoes (Mascarin et al. 2025). While sorbicillinoids from *Trichoderma longibrachiatum* SFC100166 inhibited tomato late blight, rice blast, and tomato gray mold pathogens (Ngo et al. 2021). This evidence suggests that strains under study, such as *C. rosea* TCF417 could produce sorbicillinoids relevant for plant disease suppression, including cacao diseases.

Verticillins are epipolythiodioxopiperazines (ETPs) biosynthesized via non-ribosomal peptide synthetases (NRPS). Verticillin clusters found here are widely distributed within the family *Bionectriaceae*, with *Clonostachys* spp. being the most prolific producers. They have well documented biological activities, including cytotoxic, nematocidal, antifungal, and antibacterial effects (Wang et al. 2017; Han et al. 2020; Dominique et al. 2022; Pierre et al. 2024). The BGC responsible for verticillin production was first identified in *C. rogersoniana* and comprises a single NRPS core gene (VerP), along with other tailoring genes (Wang et al. 2017; Pierre et al. 2024). In our dataset, verticillin-like BGCs were identified in multiple *Clonostachys* species, including *C. rosea* (IK726 and TCF417), *C. chloroleuca* (Cc878 and 570-77), *C. solani* 1703, and *C. rhizophaga* 906-72A. This distribution is consistent with previous reports of *Clonostachys* spp. as verticillin producers, although *C. chloroleuca, C. solani* and *C. rhizophaga* have not been previously described as sources of this metabolite (Pierre et al. 2024; Pierre et al. 2025). These clusters shared the conserved core gene and several key tailoring genes, supporting their functional similarity. Consistent with this predicted biosynthetic potential, our antifungal assays showed that verticillin D isolated from *C. rosea* TCF417 (Boya P. et al. 2024) exhibited mycelial growth inhibition against both *Moniliophthora roreri* and *Mycena citricolor*, with greater activity against *M. citricolor*. These findings provide experimental evidence of the antifungal activity of verticillin D against fungal plant pathogens. This bioactivity may explain the observed biocontrol capacity of *C. rosea* TCF417 on *M. roreri* under field condition (Castillo et al. 2025). Our findings are in agreement with previous reports of verticillin D from *C. rosea* against other biological targets, including the human parasite *Leishmania donovani* (Boya P. et al. 2024), and the fungal plant pathogens *Alternaria burnsii* and *Sclerotium rolfsii* (Dominique et al. 2022). Together, these findings support further investigation of verticillin-like BGCs and their associated metabolites as promising sources of antifungal compounds for biocontrol applications.

The hybrid NRPS–PKS cluster encoding fusaristatin A further illustrates the broad taxonomic distribution of certain biosynthetic pathways. Fusaristatin A is a cyclic lipopeptide originally isolated from the endophytic fungus *Fusarium* sp. YG-45 and its BGC was described in *Fusarium graminearum* PHI (Shiono et al. 2007 Apr 25; Sørensen et al. 2014). In our study, this BGC was identified in species from multiple genera such as in *Emericellopsis cladophorae* MUM 19.33, *Hapsidospora chrysogenum* ATCC 11550, *Waltergamsia citrina* FKII-L8-BK-P5, and the cacao endophyte *W. zeylanica* TCF400. This cluster showed conserved core genes, NRPS7 and PKS6, alongside shared tailoring enzymes, suggesting a conserved biosynthetic mechanism (Sørensen et al. 2014; Huang et al. 2024). Additionally, several genes with unknown functions were present in each cluster within the GCF. Previous studies have reported both the bioactivity of fusaristatin A and the identification of its BGC in fungal genomes. In *F. pseudograminearum*, its production was shown to reduce pathogen aggressiveness towards wheat seedlings (Khudhair et al. 2020). Also, this compound has exhibited antibacterial activity against *Xantomonas oryzae*, causal agent of bacterial blight of rice (Lim et al. 2010). The fusaristatin A BGC was identified in the genome of the marine fungus *E. cladophorae* MUM 19.33, as well as in multiple *Fusarium* species (Gonçalves et al. 2021; Lin et al. 2023). Together, this evidence supports its potential role in host-microbe interactions including endophytic associations.

A particularly notable finding of this study is the widespread presence of the squalestatin S1 BGC across all analyzed genomes. Squalestatin S1 is an inhibitor of squalene synthase and is produced by several fungi (Lebe and Cox 2019). Although squalestatin S1 has been described as a polyketide-derived compound with broad spectrum of antifungal properties (Bonsch et al. 2016), our results consistently identified a terpene-associated cluster with a conserved core gene encoding squalene/phytoene synthase. Squalene synthase (SQS) is a key enzyme in the sterol pathway and catalyzes the conversion of two molecules of farnesyl diphosphate into squalene (Linscott et al. 2016; Singh et al. 2024). Although, the core gene squalene/phytoene synthase is typically involved in primary metabolism, the observed genomic content, which includes tailoring enzymes, transporters, and accessory genes, supports its assignment as a secondary metabolite cluster. Consistent with this, similar clusters have been identified in multiple fungal genomes. For example, squalestatin S1 clusters were identified in the genomes of *Diaporthe* sp. SYSU-MS4722 (Yuan et al. 2025), *Aspergillus sydowii* MNP-2 (Fu et al. 2024) and in lichen-forming fungi containing the same core gene, squalene/phytoene synthase across several species (Singh et al. 2024). Furthermore, squalestatin S1 clusters has been identified in both biocontrol fungi and plant pathogens. For instance, it has been reported in *Talaromyces trachyspermus* IRAN 3054C, a biocontrol agent for parasitic plants affecting horticultural crops (Hemmati et al. 2026), and in *Nigrospora musae* ST1, which causes leaf spot disease in the oil-producing plant *Idesia polycarpa* (Chen et al. 2026). The broad conservation of this cluster across diverse taxa, including *Bionectriaceae*, indicates an important ecological function, possibly related to antifungal activity or competitive interactions. However, further research is needed to understand the biological activity and relevance of this terpene, against microorganisms, especially those affecting plants and whether squalestatin S1 from *C. rosea* TCF417 and *W. zeylanica* TCF400, contribute to the suppression of cacao pathogens.

In summary, the overall conserved gene organization and variation in accessory genes in several characterized GCFs indicate that *Bionectriaceae* maintain core biosynthetic frameworks while diversifying their chemical outputs. Previous studies have suggested that the high diversity of BGCs likely contribute to their fungal lifestyle, including their roles as endophytes (Kuhnert et al. 2021). Importantly, the presence of several BGC encoding bioactive SMs in cacao endophytes highlights their importance in plant health. Compounds such as sorbicillinoids and verticillins have well-documented activity against fungal pathogens, some which affect plants, reinforcing that these endophytes may contribute to the natural suppression of cacao diseases (Castillo et al. 2025). In support of this potential, our experimental assays demonstrate the antifungal activity of fractions and verticillin D from *C. rosea* TCF417 against *Moniliophthora roreri* and *Mycena citricolor*. Along with the high proportion of uncharacterized GCFs (about 75%) detected in this study, our results indicate that many potentially important metabolites remain undiscovered.

This study provides a comprehensive overview of the biosynthetic capacity of *Bionectriaceae*, including the genomes of cacao endophytes with biocontrol potential. Although the number of species studied in this work is a very small proportion of the entire family biodiversity (about 4.3% of the described species), our results suggest that the biosynthetic capacity may be greater than previously thought (Perera et al. 2023; Zhao et al. 2025). Future research would benefit from increased sequencing efforts to incorporate underrepresented species from different environments and lifestyles to validate the trends observed in this study, particularly given that most publicly available genomes belong to the genus *Clonostachys*.

## Data availability

The genome sequences generated in this study were submitted to the NCBI database under the BioProject PRJNA1169838.

## Acknowledgements

We thank Orlando Lozada owner of the farm, La Magnita, in Bocas del Toro, Panama, for his support in this research and for allowing us to collect cacao pods from which the endophytic fungi were isolated. Also, to Ana Gabriela Moreno and Eunice Tapia, for their technical support with laboratory work; and to the MIAMBIENTE granting the specimen-collecting permits to do this research.

## Study funding

Funding for this research was provided by the National Secretariat for Science, Innovation, and Technology of Panama (SENACYT) (grant numbers FID17-104 and FIED25-20), and a grant from the Ministry of the Environment of Panama (Biobank Project MIAMBIENTE UFISI-003-49-2019). Financial support was additionally provided by the Global Environmental Facility (GEF-UNDP, Project PS97410). L. C. Mejía and M. Gutierrez also acknowledge support from the SENACYT SNI program.

## Conflicts of interests

The authors declare no conflicts of interests.

## Author contributions

LAA, MG, and LCM contributed to the conceptualization and design of the study, as well as to the funding acquisition. LAA, HEC, MBJ, IQ, AM, and CR performed the experiments and material preparation. Data collection and analysis were performed by the LAA. The first draft of the manuscript was written by LAA and LCM. LAA, HEC, MBJ, IQ, AM, CR, MG, and LCM have reviewed and commented on previous versions of the manuscript. All the authors have read and approved the final manuscript.

## Supplementary material

Figure S1. Squalestatin S1 gene cluster family (GCF) synteny analysis.

Figure S2. Temporal dynamics and treatment comparisons of mycelial growth inhibition (GI%) by fractions from *Clonostachys rosea* TCF417 against *Moniliophthora roreri* and *Mycena citricolor*.

Figure S3. Antifungal activity of fractions (F1-F4) and verticillin D from *C. rosea* TCF417 against *Moniliophthora roreri* and *Mycena citricolor* at 13 days post-inoculation.

Table S1. Fungal strains and associated genomes from *Bionectriaceae* family analyzed in this study.

Table S2. BUSCO completness assessment (assembly mode) for the genomes of the fungal isolates *Clonostachys rosea* TCF417 and *Waltergamsia zeylanica* TCF400.

Table S3. Genome assembly and annotation statistics of *Clonostachys rosea* TCF417 and *Waltergamsia zeylanica* TCF400.

Table S4. Distribution of Biosynthetic Gene Cluster (BGC) categories identified in 32 fungal genomes of the *Bionectriaceae* family using antiSMASH.

Table S5. Gene cluster families (GCFs) identified by BiG-SCAPE v2.0 at a distance cutoff of 0.5, based on antiSMASH-predicted BGC regions from 32 fungal genomes analyzed together with MIBiG 3.1 reference clusters.

Table S6. Biosynthetic gene cluster families (GCFs) in *Bionectriaceae* with characterized BGCs using BiG-SCAPE at cutoff 0.5.

Table S7. Summary of Analysis of Variance (ANOVA) for the growth inhibition (GI%) of *Moniliophthora roreri* and *Mycena citricolor* exposed to fractions (F1-F4) and verticillin D obtained from *C. rosea* TCF417 at 7 and 13 days post-inoculation.

Table S8. Post-hoc Tukey HSD multiple comparison test for the growth inhibition (GI%) of *Moniliophthora roreri* and *Mycena citricolor* treated with fractions (F1–F4) obtained from *C. rosea* TCF417 at Days 7 and 13 post-inoculation.

Table S9. Temporal dynamics of growth inhibition (GI%) for *Moniliophthora roreri* and *Mycena citricolor* treated with verticillin D and Mancozeb across evaluation days (Days 3 to 13).

Table S10. Post-hoc Tukey HSD multiple comparison test for growth inhibition (GI%) of *Moniliophthora roreri* and *Mycena citricolor* treated with verticillin D at days 7 and 13 post-inoculation.

